# Fear-Mouse Tracker (FMT): An open-source toolkit to track innate defensive behaviors with high precision

**DOI:** 10.1101/2022.11.25.517925

**Authors:** Sanket Garg, Gabriela Pino, Claudio Acuna

## Abstract

In the past years, machine-learning-based approaches to track animal poses with high spatial and temporal resolution have become available, but toolkits to extract, integrate, and analyze coordinate datasets in a user-friendly manner have lagged behind. Here, we introduce Fear-Mouse Tracker (FMT), a simple and open-source MATLAB-based pipeline to extract and quantitatively analyze DeepLabCut-derived coordinates of mice presented with threatening stimuli that commonly trigger innate defensive responses. This framework allows for unbiased quantitative estimations of stretch-attend posture (SAP) observed during risk assessment behaviors, as well as for measurements of the timing and extent of freezing and escape responses that follow the presentation of threatening stimuli such as a predator odor, or sweeping and looming stimuli resembling predator approaches. FMT is specially designed for users not very experienced in using programming languages, thus making it more accessible to a broader audience.

## INTRODUCTION

Innate defensive responses are a group of evolutionary conserved behaviors, commonly triggered by threatening stimuli^1-8^. These responses typically include risk assessment, flying, freezing, an in many cases defensive attacks. Innate defensive behaviours are controlled by a collection of brain structures including the amygdala^9^, the hypothalamus^10-15^, and the midbrain^9,10,16,17^. The midbrain’s Periqueductal Gray (PAG) is the final common path for all types of defensive responses^16,17^, and thus unraveling how PAG circuits and cell types promote and regulate specific aspects of defensive responses represents a major research challenge in neuroscience. Importantly, deficits in defensive responses are often linked to anxiety disorders, the most prevalent mental disorders of our time.

To understand how exactly the PAG promotes and regulates defensive behaviors, it is crucial to be able to manipulate specific PAG neural pathways and cell types in awake behaving mice, to acquire rich data representations of these behaviors, and to be capable of dissecting their dynamics quantitatively. Over the last decade, mostly with the development of tools to gain genetic access to specific cells and circuits^18^ and with the rise of chemo- and optogenetic technologies^19,20^, assessing the contribution of subsets of cells and circuits to specific behavioral outputs has become possible. However, development of tools to quantitatively assess naturalistic behaviors with high spatial and temporal resolution has lagged behind^21^.

Here, we implemented the machine-learning based open-source toolbox DeepLabCut (DLC)^22,23^ to track mouse movements and poses upon presentation of threatening stimuli that trigger innate defensive responses, and then developed a MATLAB-based toolkit to efficiently extract, integrate, and quantify the dynamics of risk assessment as well as flying and freezing responses. We expect this framework will greatly facilitate and accelerate dissecting the roles of different PAG cell types, neural circuits, and synaptic mechanisms in promoting and regulating defensive behavior in normal mice as well as in mouse models of anxiety disorders.

## MATERIALS AND METHODS

### Mice

12-week-old male animals (C57BL6/J background) were used in all experiments. Mice were maintained on a 12-hour light/dark (7pm/7am) cycle, with ad libitum supply of food and water. All behavioural tests were run in the animal’s active period. All animal studies were approved by the Governmental Council Karlsruhe, Germany.

### Predator-odor Test

The test was performed in the animal’s own cage, under dim red light (55 LUX). The stimulus threat consisted of a fabric with cat odor, which was obtained by petting a cat with a fabric for 2 weeks (1 daily session of 30 minutes), and then stored at -20°C. Animals were habituated for 4 consecutive days (1 daily session), and the test was run on the 5^th^ day. Mouse behavior was simultaneously recorded with a top- and a side-view camera (C920 Webcam, Logitech), using AnyMaze (Stoeling Co.). Each habituation session consisted of exposing the mice to 3-4 small pieces of control fabric (identical to the one with the cat odor, except that it did not have the smell of the cat) wrapped into knots. Fabrics were placed in the lower left corner of the cage (as shown in Fig. 2A) and animal’s behavior was monitored for 10 minutes. On the test day, a recording of the basal activity of the animal towards the control fabric was done for 5 minutes. After this, the control fabric was replaced by the cat odor fabric, and the animal’s behaviour was recorded for 10 minutes.

### Looming and Sweeping Tests

These tests were performed in a red plexiglass open field arena (50×50×35cm, 20 LUX), comprising a small sliding door and a mouse shelter. Mice were video-tracked with 3 IR USB night vision cameras (ELP, Shenzhen Ailipu Technology) located on top, and on two opposite sides of the arena, controlled by AnyMaze (Stoeling Co.). Videos were recorded at 30FPS, 1080p. Looming and sweeping visual stimuli were generated by a custom-written Python script. Looming stimuli consisted of a black circle, presented at the center of the screen, expanding from 5 to 25cm of radius at 45cm/s. Sweeping stimuli consisted of a black circle moving diagonally from the upper corner to the opposite bottom corner of the screen. In both loom and sweep experiments, the contrast (looming test, 10-100%) or the size and speed (sweeping test, 1.5 to 4cm radius and 4 to 10 cm/s, respectively) of the stimulus was varied throughout the test to prevent habituation and to modulate the vigour of the evoked defensive responses. Visual stimuli were presented via an LCD PC-screen (Asus, Japan) held on top of the arena, covering nearly half of the arena area (see Fig.3). The shelter was placed at one of the arena’s corner, outside the area covered by the LCD screen. **Habituation sessions**. Mice were habituated to the arena for 4 consecutive days, and the test was run on the 5^th^ day. During habituation, mice were allowed to enter the arena by the sliding door, and then freely explore it for 10 minutes. **Test session**. On the test day, mice were introduced into the arena as for habituation days, and then presented with either looming or sweeping stimuli, whenever the animal was located under the top screen and far away from the walls. Each stimulus was presented at least 10 times to each mouse, at inter-stim intervals of at least 1 minute. Behaviors were continuously recorded with all cameras during the whole test duration.

### Extracting mouse coordinates using DeepLabCut

Deeplabcut (DLC)^22,23^ was used to extract mouse coordinate during defensive responses. For this, we created two separate networks, one for the cat odor experiments and another one for loom-sweep experiments, both based on top-view video-recordings. DLC uses a kmeans algorithm to cluster the frames based on the visual appearance and then to extract a set number of frames from each video. For the cat odor and loom-sweep experiments 650 and 200 frames were extracted, respectively. These frames were then manually labeled. For all the videos, 4 markers were used to define the boundaries of the experimental setup. In addition, for cat odor experiments, 9 markers were placed along the length of the animal starting from the nose till the tailbase, along with markers for the left and right ears. For looming/sweeping experiments, three markers were placed on the mouse body (i.e., nose, bodycenter and tailbase) and an additional marker was also placed at the bottom, top-left corner of the rectangular shelter. A percentage (0.05) of the complete labeled dataset was then set aside to create the test set and the remaining dataset (0.95) was used for training the resnet_50 network, for a total of 250,000 iterations. The network was evaluated at steps of 10,000 iterations to determine how the training and test errors reduced with increasing iterations. All the videos were analyzed using the trained network, outlier frames were extracted using the ‘jump’ algorithm, and then manually corrected, resulting in a total of around 1100 frames for the cat odor videos and 600 frames for the loom-sweep videos following which the network was retrained for another 250,000 iterations. The snapshot at 250,000 iterations was used for generating mouse posture coordinates from all the cat odor videos, while the snapshot at 240,000 iterations was used for generating coordinates from all the loom-sweep videos. The train and test errors achieved at these iterations closely matched human accuracy levels of nearly 2.5 pixels (Fig. 1).

**Figure 1.**
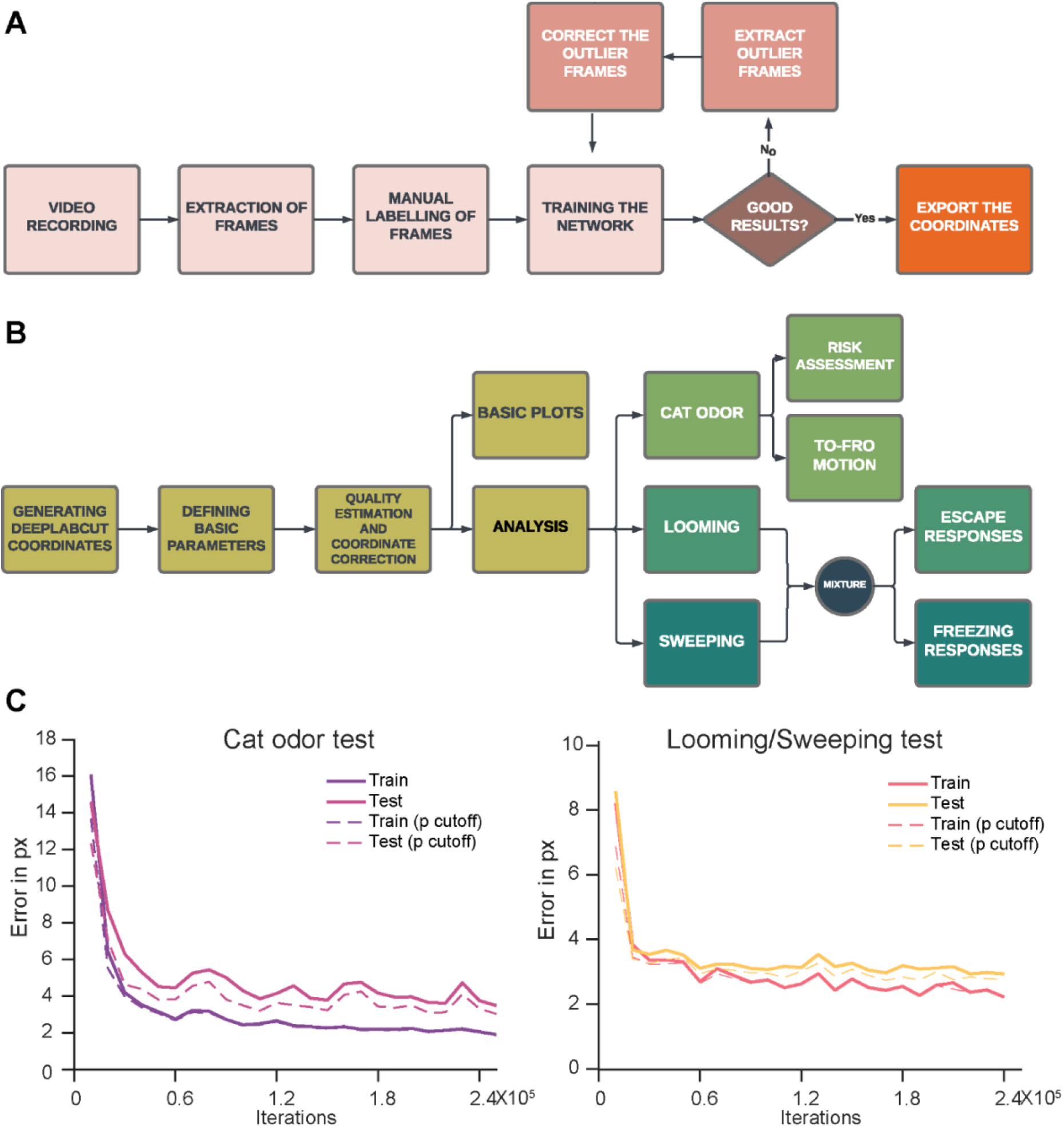
Fear-Mouse Tracker (FMT): A framework for quantitative assessment of innate defensive behaviors in mice. **A**. Flowchart depicting DLC^23^-based extraction of mouse coordinates. Video-recordings are used to first extract frames, which are then manually labeled and used for training the network. The trained network is then used to extract mouse coordinates. If the results are good, then coordinates are exported, else outlier frames are discarded, corrected, and the network is retrained. **B**. Flowchart depicting the FMT framework we developed. Basic parameters are defined for analysis. DLC coordinates are subjected to quality control and correction, and then used for behavioral analysis. In cat-odor experiments, risk assessment and To-fro is measured, while properties (timing and extent) of escape and freezing responses are analyzed in loom-sweep experiments. **C**. Left. Test and train error versus number of iterations for the cat odor experimental conditions. **D**. Test and train error versus number of iterations for the loom and sweep experimental conditions. The p cutoff used for the dotted plots in figures **C** and **D** was 0.6.

### Analysis of cat-odor experiments

The coordinates obtained from Deeplabcut were all in pixels. In order to analyze the mouse length, distance traveled, and velocity, and to allow for comparison across videos, it was necessary to convert these pixel coordinates to real world centimeter-based coordinates. In the case of the cat odor experiments, the length and the breadth of the experimental box had been manually noted to be 36.3 cm and 15.4 cm respectively, and these were used to generate the pixel to centimeter ratio for conversion to real world coordinates. This pixel-to-centimeter ratio was used to convert all the 11 body markers on the mouse and the 4 field markers to real world coordinates. These corrected markers were then used for analyzing SAP and to-fro behavior across the field.

### Analysis of loom-sweep experiments

In the case of loom-sweep videos, the top camera had to be placed at a 10-25 degree angle, owing to the presence of the LCD monitor required to project visual stimuli (see Fig. 3). As a result, the square experimental field now appeared like a trapezium. The experimental area’s dimensions were 50cm X 50cm, and the camera was placed at the height of 35 cm. With this information, a geometric approach was used to calculate the actual real-world coordinates of any marker. To calculate the x-coordinate, we used a ratio-proportion-based method by analyzing how the front edge changed to the back edge in terms of the pixel lengths. For the y-coordinate, we first used a projection method to compute what angle of inclination, given the manually measured range, could convert a square to the current trapezium. This angle was then used to measure the distance of any marker from the front edge. We then created a new variable called the location variable (*loc*), to account for inaccurate location estimations observed when the mouse was located within the shelter, or in the field but close to the edges or around the corners. This *location* variable was derived from the transformed bodycenter, nose, and tailbase markers, in that order, taking into consideration the likelihood values which DLC had provided. For every frame, the *loc* took the value of the bodycenter coordinate if its likelihood value was greater than 0.9, the nose coordinate if its likelihood value was greater than 0.9, or the tailbase coordinate if that acquired a likelihood value greater than 0.9. In some cases, some frames (less than 1% of total frames in all sessions, except for 3 sessions in which the values were greater than 1%.) were left that did not receive any value. The best estimate for the location of the mice in these frames was computed by constructing a straight line between the last and first existing values around those frames. *Loc* was then utilized to quantify freezing, escape, or a mixture of the responses triggered by looming or sweeping stimuli. **Escape Responses**. We defined an escape response as the quick movement of the animal into the shelter, and this response must be time-locked to the stimulus presentation. The escape response was classified as a fast escape response if the maximum speed exceeded a threshold (40 cm/sec), or else a slow escape. If the animal only succeeded to reach close to the shelter, then the response was classified as an attempted escape. The beginning of the escape response was defined as the moment when the animal’s velocity increased above 5 cm/sec, towards the maximum value and the end was defined as the moment the animal entered the shelter. In order to track the animal’s position with respect to the shelter, we made use of the top-left corner of the shelter’s base (see mark in Fig. 3A). The distance between *loc* and the shelter coordinate was computed for every frame. This distance value was negative if the animal was present within the shelter, and positive if the animal was present outside the shelter. **Freeze Responses**. Freezing behavior was defined as the periods whenever the animal remained stationary for some duration. The freeze detection algorithm allowed for the identification of all these periods. The algorithm is based on creating a small square around the location of the animal for every frame, and calculating for how many frames, the animal stays within the square defined by the current frame. Our approach allows to calculate the exact beginning and end of each of these freezing periods. The freezing responses are classified as being long freeze behavior if the freezing duration is greater than a second for at least one of these periods. Else, if the freezing duration is greater than 300ms for at least one of these periods, the behavior is classified as a short freeze.

### Codes

The FMT MATLAB codes can be accessed at the following Github link: https://github.com/AcunaLabUHD/FMT.

## RESULTS

### The pipeline

To achieve accurate coordinates for our analysis, we created two independent DLC^22,23^ networks, one for the cat-odor experiments and another one for the looming-sweeping experiments. These networks were created by extracting frames from all the videos and labeling a few (650 and 200 for cat-odor and sweep-loom experiments, respectively), which were then used to train the networks. If the results were not satisfactory, outlier frames were extracted, manually corrected, and included as part of the training data. The pipeline for the DLC training is summarized in Fig 1A. During the training iterations, train and test errors were recorded at intervals of 10,000 iterations, and the training process was continued till train and test errors were reduced to values in the range of 2.5-3 pixels, resembling human-level accuracy (Fig 1C). The resultant networks were then used to analyze all experimental videos. Specifically, the coordinates generated were fed into the FMT algorithm (Fig 1B). FMT begins with defining a few basic parameters like the frame rate, file directory paths, and other manually changeable variables. The coordinates are first analyzed for quality measures based on the likelihood values, are subjected to distance unit transformations and corrections (trapezium to square correction for loom-sweep experiments), and then used for analysis. FMT measures quantitatively risk assessment-induced stretch-attend posture (SAP)^24^ behavior and To-fro motion (including escape responses) across the length of the experimental field in cat odor experiments, as well as the timing and extent of escape and freezing behavior in sweeping/looming experiments.

### Cat-odor experiments

We first tested the ability of FMT to extract behavioral events in mice presented with cat-odor that triggers prominent defensive responses (Fig. 2A, B). For analysis of these behaviors, boundaries were created at 25% and 75% of the length of the mouse cage, defining a danger zone and a safe zone (Fig. 2A). The fabric with (test) and without (control) cat odor was placed on the lower corner of the danger zone. Mice exposed to fabrics containing cat odors (but not those exposed to control fabrics) spend more time in close proximity to the fabrics, as revealed by density plots (Fig 2C), display risk-assessment behaviors (Fig. 2D, E), To-fro motion between zones, and escapes responses into the safe zone (Fig. 2F, G). The Main_CatOdor.m and the Risk_function.m are the FMT functions that perform such analysis. **Risk-assessment**. We measured risk assessment by quantifying SAPs^24^. For this, the length of the animal was computed in each frame by summing the individual distances between all the 9 markers placed along the antero-posterior axis of the mouse. The basal videos were used to compute the average and the standard deviation of the animal length in the period starting from placing the fabric (with no odor) until 3 minutes later. An upper threshold was defined as the sum of this mean and standard deviation. The nose of the animal was used to define its position in the danger, middle or safe zone, and the average length in the three zones was computed along with the average length for the entire analysis period. Besides, the average length above the upper threshold and the average length above the upper threshold in the danger zone were also computed. SAP was defined as frames in which the animal’s length crosses the upper threshold. This was used to calculate the frequency and duration of SAP behavior in the danger zone and in the whole arena. **Escape responses**. In order to estimate escape responses, the movement of mice between the danger and the safe zones was assessed. The bodycenter mark defined the animal’s presence in either of the three zones. The frames at which the animal left the danger zone, entered the safe zone, left the safe zone, and re-entered the danger zone were identified. The movement of the animal from the danger to the safe zone was defined as an escape response, and the movement of the animal from the safe to the danger zone was defined as an approach response. The paths of the animal during these escape responses, their average escape velocities, and the distance covered were compared in the basal and the cat odor conditions. **To-fro**. Each instance when the animal left the danger zone, travelled to the safe zone and returned to the danger zone was defined as a single to-fro behavior. The number of such To-fro responses were computed in the period till the animal removed the fabric from the danger zone. The number of To-fro responses as a fraction of the total frames, starting from placing the fabric till the animal removed the fabric from the danger zone, was then calculated.

**Figure 2.**
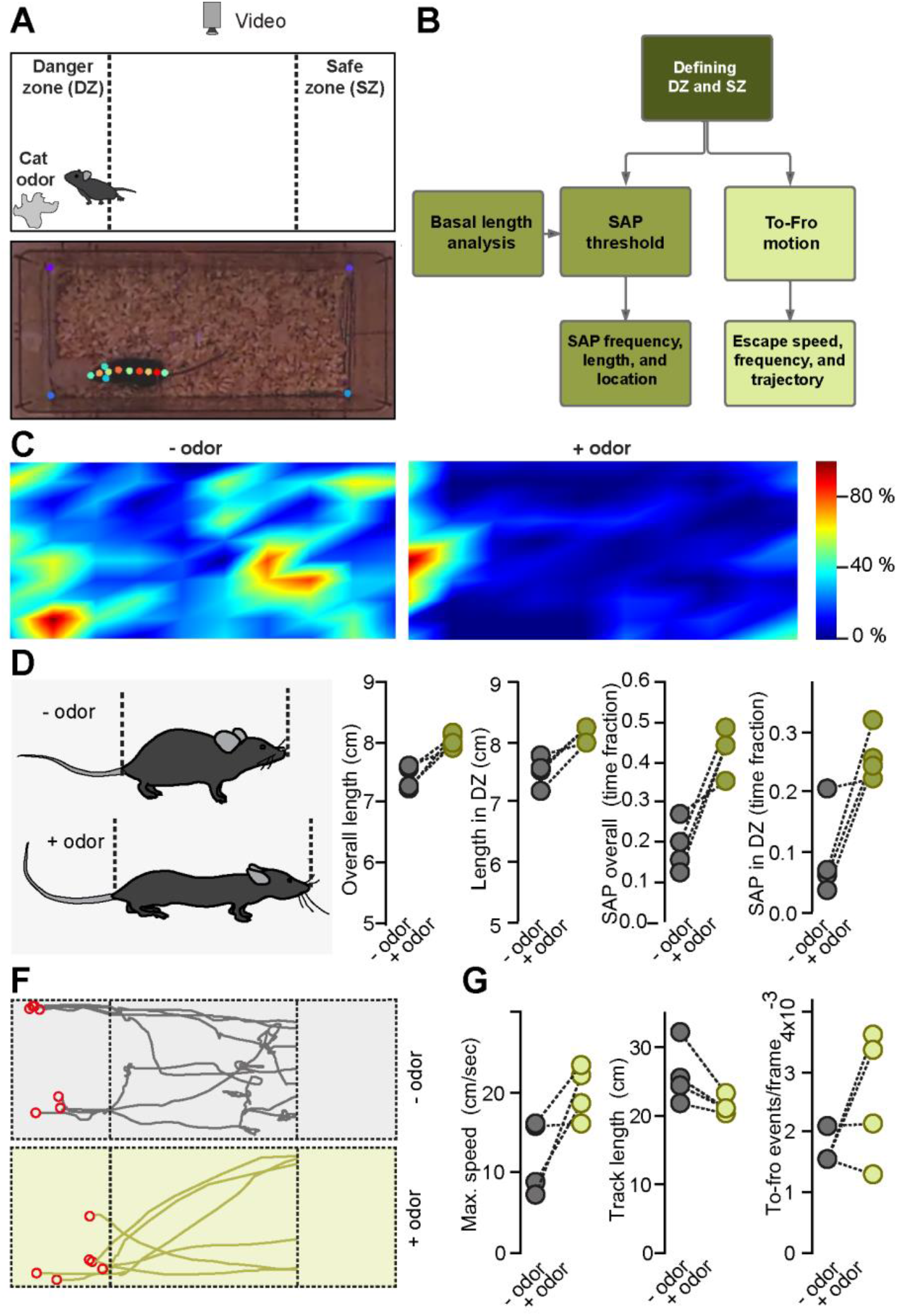
Using FMT to extract behavioral events triggered by predator odor. **A**. top, schematic of experimental configuration. Bottom. Marks used to track mouse movements and postures with high accuracy via DeepLabCut (DLC). **B**. Flowchart showing features and workflow used by FMT to extract behavioral events derived from cat-odor experiments. **C**. Density plot displaying mouse location before and after exposure to a cat odor. The fabric containing cat odor was placed on the lower left corner as shown in **A. D**. Schematic of postures shown before and after exposure to cat-odor. This posture is called stretch-attend posture (SAP), and is commonly observed during risk assessment behaviors. **E**. Summary graphs of mouse length in different zones of the cage, and total fraction of time used fby a mouse in SAP before and after presentation of Cat odor. **F, G**. Tracks of mouse trajectories (from the danger zone into the safe zone) before and after exposure to cat odor. Left, representative escape responses. Red circles highlight the onset of a behavioral event. Right. Summary plots displaying properties of escape responses.

A representative experiment and summary graphs showing averaged behaviors in 4 mice is presented in Fig. 2C-E. Typically, mice exposed to anaesthetized predators, or predator odor explore significantly the predator or the source of predator smell (Fig. C), perform risk assessment behaviors (Fig. D, E), and then execute motor response that typically comprise escape events (Fig. F, G). FMT proved capable of reliably extracting each of these behavioral events. For instance, risk assessment behaviors typically comprised stretch-attend postures (SAP), characterized by increases in the length of the animal’s body. As shown in Fig. 2C, D, FMT can reliably track the timing, frequency, and extent of SAP events, and relate them to particular parts of the arena such as the danger zone. Last, motor behaviors triggered by presentation with threatening odors normally include escape responses. FMT could reliably identify those responses, and accurately track their frequency, speed, directionality (track-length). Altogether, these experiments indicate that FMT can reliably track and quantify specific components of defensive responses triggered by naturalistic stimuli such as the presence of a predator odor.

### Loom-sweep experiments

Laboratory mice display prominent innate defensive behavior when presented with looming and sweeping stimuli. Typically, a looming stimulus consists of a high contract shape that enlarges at a given speed. This resembles an approaching predator, which produces looming shadows. In contrast visual sweeping stimuli consist of a high-contrast shape that does not change in size but moves across the screen at a particular constant speed, resembling a predator cruising in the sky. Both visual stimulation protocols trigger defensive responses reliably, and thus are widely used to uncover the neural basis of innate fear. We therefore set out to implement FMT in the context of these two behavioral paradigms (Fig 3). **Analysis window**. This is defined by *curr_duration*. Looming stimuli typically last a few hundred milliseconds (in our conditions, around 500 ms) and trigger behavioural events for several seconds. Therefore, FMT uses a period of 8 seconds, starting from the onset of the stimulus, as analysis period. In contrast, sweeping stimuli lasts for at least a few seconds (normally more than 5 seconds), and thus we chose the period between the start and end of the stimulus as the period for analysis. **Escape responses**. After observing the animal’s behavior post-stimulus presentation, we noticed that in most looming trials and some sweeping trials, the animal has a quick escape response to the shelter, often reaching it in less than 2.5 seconds of stimulus onset. FMT uses two conditions to define escape responses. First, escape responses require the animal to reach the shelter within 2.5 second after stimulus presentation. This parameter is defined in *time_fastEscape*. Second, the maximal speed reached during the escape responses allow classifying them as fast (>=40 cm/sec) or slow (<40 cm/sec) escape responses. This value was chosen by observing the range of velocities of the animal in response to various stimuli. **Freezing responses**. Freezing is commonly triggered by sweeping stimuli, and to a lesser extent by looming stimuli. To extract freezing events, we computed the maximum length stationarity (see Methods for details) depicted by the animal after the stimulus onset out. We classified the freezing responses as being a long freeze if this maximum duration exceeded a second, a short freeze if it exceeded 300 ms, and no freeze for other cases. The reason for such a time-based classification of freezing responses was that we wanted to differentiate any brief freezing responses the animal showed just before an escape response from the longer freezing responses where the animal eventually may or may not have shown an escape response. **Mixed escape-freezing responses**. In many cases, loom-sweep stimuli triggered a mixture of freezing and escape responses. This normally occurs when the animal does not develop stimulus-locked escape responses. In such a scenario, the animal can or cannot reach the shelter within *curr_duration*. If the animal reaches the shelter within *curr_duration*, FMT checks for freezing behaviors and classifies them as described above. Following freezing events, FMT runs again the escape detection algorithm described above and classifies escape responses according to the *maxspeed_fastEscape* threshold. If the animal does not return to the shelter by the end of the *curr_duration* due to very long freezing responses, FMT checks for any escape response to the shelter within the *time_fastEscape* period after freezing. **Attempted escapes**. These rare events represent approaches to the shelter (within 5 cm) in the *time_fastEscape* period (2.5 seconds), without entering it.

**Figure 3.**
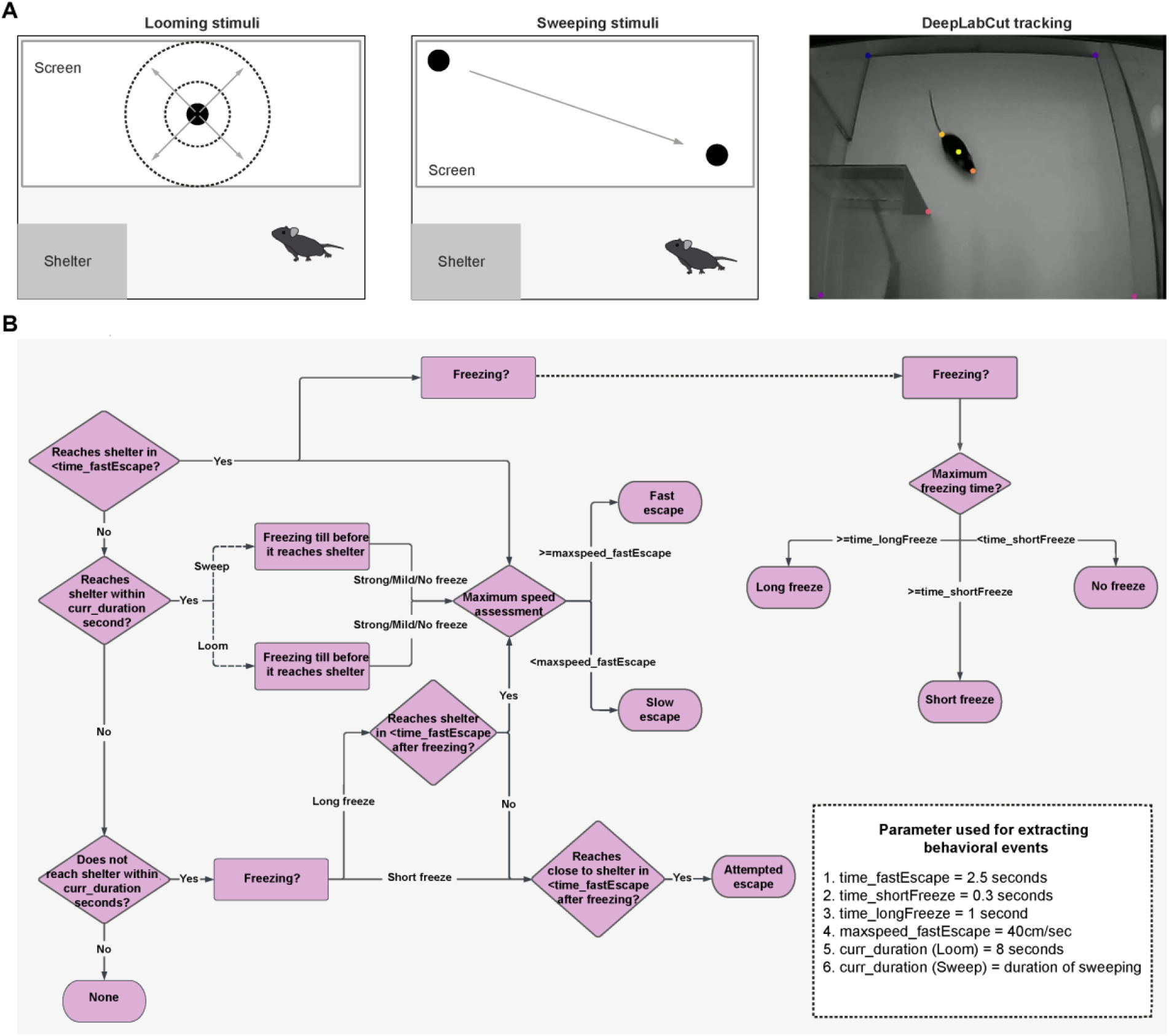
FMT workflow to extract behavioral events in looming and sweeping experiments. **A**. top and middle: schematic of experimental setup used for looming (top) and sweeping (middle) experiments. Bottom: top-view image of a typical experiment, highlighting a mouse and the marks used to track movements and postures using DLC. **B**. Flowchart representing the pipeline followed to classify behavioral responses as freezing, escape or a mixture of both. The different parameters defined during the process are mentioned at the bottom right.

We used the analysis pipeline described above to automatically and quantitatively analyse mouse defensive responses upon presentation of looming (Fig. 4) and sweeping (Fig. 5) visual stimuli. **Looming experiments**. Looming stimuli consisted of a round black circle on a white background, that grew progressively larger at a fix speed (45cm/s) but with different contrasts (10-100%). Exemplary traces of escape responses triggered by maximal looming stimuli (100% contrast), as well as of the timing and speed of such behaviors in one typical mouse, is shown in Figure 4 A, B. Typically, and consistent with previous findings^25,26^, maximal loom stimuli consistently triggered short-delay (average delay: 0.25±0.05 sec, mean±SEM) and high-speed escape responses (120.1±8.4 cm/sec, mean±SEM) in most trials (Fig. 4C, D), consistent with previous reports. In contrast, low-contrast looming stimuli (<100% contrast) triggered mixed responses comprising pure escape, pure freezing, or a mix of both behaviours (Fig. 4C, D). A summary of the delay, max speed of escape, and duration of freezing for all 7 mice analysed, segregated by stimulus parameter is shown in Fig. 4D. **Sweeping experiments**. Six mice were presented with sweeping stimuli, which vary in size and speed (1.5-4cm radius and 4-10 cm/s), and then FMT was used to quantitatively analyse their defensive responses. In contrast to looming stimulus, sweeping stimuli prominently triggered freezing responses as reported previously^27^ (Fig. 5A, B). The extent and timing of sweeping-evoked responses varied according to the speed of stimuli (Fig 5C, D): sweep stimuli passing by at speeds larger than 10cm/sec routinely triggered mixed responses composed of both freezing and escape responses, while lower speed stimuli trigger mostly freezing responses (Figure 5C, D). A summary of the delay, max speed of escape, and duration of freezing for all 6 mice presented with sweeping stimulation of different speeds, is shown in Fig. 4D. Taking together, these proof-of-principle experiments indicate that FMT can reliably track, dissect, and analyse defensive responses triggered by looming and sweeping stimulation, which are commonly used to study the neural basis of innate defensive behaviors in laboratory mice.

**Figure 4.**
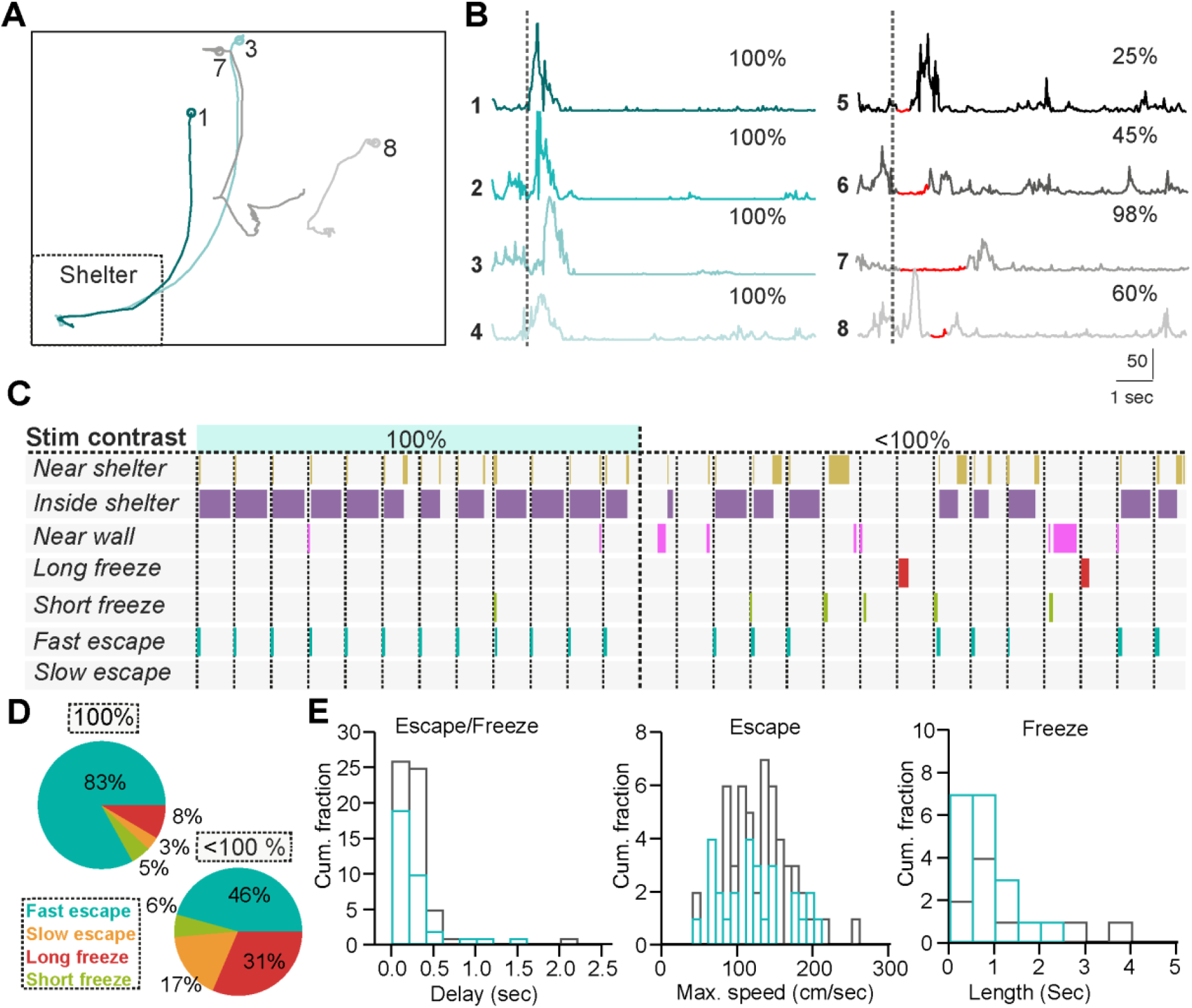
FMT-mediated dissection of defensive responses triggered by looming stimuli. **A**. Representative behavioral responses triggered by looming stimuli of different contrasts (10-100%). **B**. Plots of mouse speed as a function of time before and after presentation of looming stimuli (vertical dotted line). High-contrast looming stimuli (100%) triggered mainly escape responses (upward deflections), while low-contrast stimuli triggered freezing events (red), typically accompanied of escape responses (upward deflections). **C**. Timeline of looming-evoked behavioral events categorized using FMT. Vertical dotted lines represent presentation of looming stimuli in individual trials. **D**. Summary pie charts showing categorization of responses triggered by looming stimuli of different contrasts in all experiments. Data derived from 7 animals was pooled. **E**. Histograms summarizing delay, max speed of escape, and duration of freezing in 7 mice (72 trials in total). Grey: 100% contrast. Blue: <100% contrast.

**Figure 5.**
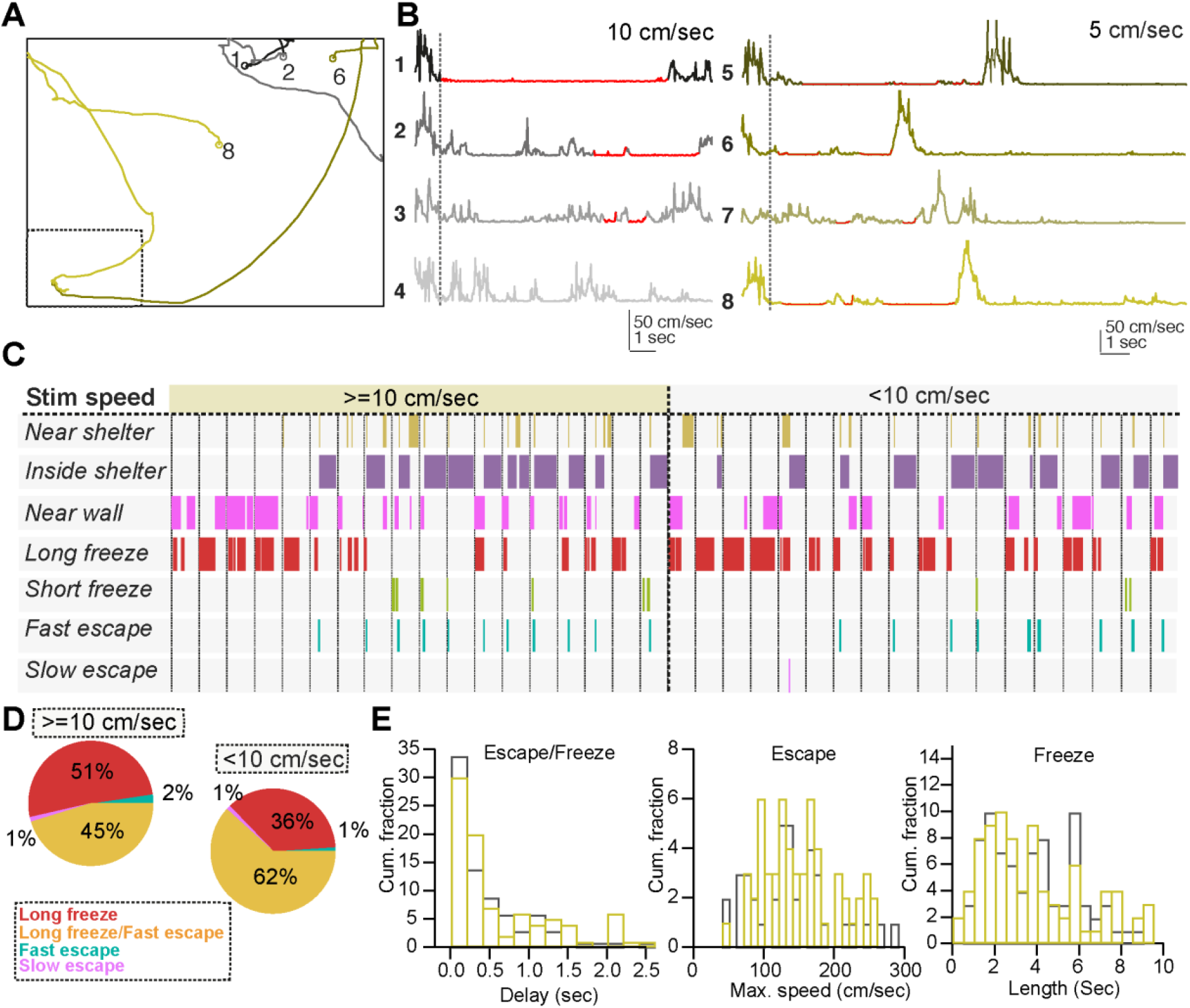
FMT-mediated dissection of defensive responses triggered by sweeping stimuli. **A**. Representative behavioral responses triggered by sweeping stimuli of different speeds (2-15 cm/sec). **B**. Plots of mouse speed as a function of time before and after presentation of sweeping stimuli (vertical dotted line). High-speed sweeping stimuli (10-15 cm/sec) triggered mainly freezing responses (red), while low-speed stimuli triggered mixed freezing/flying (up to 10 cm/sec, red/upward reflections) responses. **C**. Timeline of sweeping-evoked behavioral events categorized using FMT. Vertical dotted lines represent presentation of sweeping stimuli in individual trials. **D**. Summary pie charts showing the overall (population) categorization of responses triggered by sweeping stimuli. Data derived from 6 animals was pooled. **E**. Histograms summarizing delay, max speed of escape, and duration of freezing in 6 mice (107 trials in total). Grey: 10-15 cm/sec speeds. Green: 5-10 cm/sec speeds.

## DISCUSSION

Over the last decade, a large number of new methods to trace^28,29,30^, manipulate^19,20^, and watch^31,32^ the activity of neural circuits has emerged. These technologies have greatly helped establishing causal links between neural activity of specific cell populations in the brain and behavioral outputs. Parallel development of methods to assess behavioral outputs quantitatively has lagged behind^21^, but with the recent emergence of machine learning based approaches to track animal’s movements and postures with high precision, this has begun to change^22, 23^. In this paper, we describe a simple and open-source MATLAB-based toolkit to quantitatively assess defensive behaviors in laboratory mice. Our framework, which we called FMT, is expected to facilitate identifying and quantifying behavioral motifs underlying defensive behaviors and to help linking, quantitatively, neural activity patterns to specific behavioral events.

In the context of defensive responses, behavioral scoring in most early studies relied on human observers^33-35^, which is time consuming, can be subjective, and therefore is often prone to errors. More recent studies have used a combination of manual and automatized scoring of defensive behaviors, but these methods often lack flexibility for categorizing freezing or escape responses, and set manual thresholds for detecting specific behavioral events^4,25-27,36-39^. The analysis pipeline developed here helps to solve these problems, and thus is expected to greatly facilitate automatic, precise, and quantitative estimations of behavioral events commonly observed upon presentation of threatening stimuli to laboratory mice. First, it relays on mouse coordinates obtained via DeepLabCut^22,23^, which tracks user-defined animals’ body-parts with high accuracy and speed (Fig. 1). Second, it can accurately quantify the frequency, dynamics, and extent of risk assessment behaviors that occur prior to completion of freezing or escape in animals exposed to natural predators such as a rat or to odors of natural predators such as cat-odor. For this, FMT computes mouse’s body length in each frame by summing the individual distances between all 9 markers placed along the antero-posterior axis of the mouse (Fig. 2). Third, FMT can accurately dissect the behavioral repertoire triggered by presentation of looming and sweeping visual stimuli (Figs. 3-5), commonly used to study the neural basis of innate fear in laboratory mice. Specifically, it can extract and classify the timing and vigor of escape and freezing responses based on accurate estimations of mouse speed, stationarity, and movement trajectory. Fourth, it is inexpensive and user-friendly. These are important factors as neuroscience is becoming increasingly multidisciplinary, highlighting the need to make behavioral analysis tools accessible to users not very experienced in programing languages.

While FMT already represents an important step towards studying defensive behaviors quantitatively, further developments aiming to improve the identification of stereotyped postures and movements are warranted. Along this line, future efforts will aim to extend this framework to other more naturalistic behavioral tests used to study innate fear, such as the rat exposure test^11^, and to implement three-dimensional tracking of mice undergoing defensive responses, likely via triangulation of multiple video-signals^40^ and depth-sensing video-tracking^41,42^. This will be incredibly useful, for instance, to dissect risk assessment behaviors, that encompass not only elongation of animal’s body but are also accompanied of changes in the positioning and directionally of limbs, tail, and whiskers, and more generally in their three-dimensional postures upon presentation of a highly threatening stimuli.

Taken together, the development of quantitative frameworks to assess defensive responses such as FMT are expected to greatly enhance our ability to deconstruct the microstructure of these behaviors and to determine how specific circuits, cell types, and synaptic mechanisms encode specific behavioral events, not only in the normal brain but also in disease models in which defensive behaviors are dysregulated. Along this line, we expect FMT will help identify abnormalities in mouse models of anxiety disorders, and importantly, in assessing the impact of novel drugs aiming to correct these behavioral deficiencies. Thus, FMT might be instrumental in accelerating our quest to find better drugs to treat anxiety disorders that represent an immense burden for society worldwide.

